# Altered Cell-Substrate Behavior on Microporous Membranes is a Result of Disruption and Grip

**DOI:** 10.1101/563361

**Authors:** Zahra Allahyari, Shayan Gholizadeh, Henry H. Chung, Luis F. Delgadillo, Thomas R. Gaborski

**Affiliations:** Department of Microsystems Engineering, Rochester Institute of Technology, 160 Lomb Memorial Drive, Rochester, NY 14623, USA; Department of Biomedical Engineering, Rochester Institute of Technology, 160 Lomb Memorial Drive, Rochester, NY 14623, USA; Department of Biomedical Engineering, University of Rochester, 201 Robert B. Goergen Hall, Rochester, NY 14627, USA

**Keywords:** Cell spreading, cell migration, fibronectin, polyethylene glycol, porous membrane, cell-substrate interactions

## Abstract

Porous membranes are ubiquitous in cell co-culture and tissue-on-a-chip studies. These materials are predominantly chosen for their semi-permeable and size exclusion properties to restrict or permit transmigration and cell-cell communication. However, previous studies have shown pore size, spacing and orientation affect cell behavior including extracellular matrix production and migration. The mechanism behind this behavior is not fully understood. In this study, we fabricated micropatterned non-fouling polyethylene glycol (PEG) islands to mimic pores in order to decouple the effect of surface discontinuity from grip provided by pore wall edges. Similar to porous membranes, we found that the PEG islands hindered fibronectin fibrillogenesis with cells on patterned substrates producing shorter fibrils. Additionally, cell migration speed over micropatterned PEG islands was greater than unpatterned controls, suggesting that disruption of cell-substrate interactions by PEG islands promoted a more dynamic and migratory behavior, similarly to cells migrating on microporous membranes. Preferred cellular directionality during migration was nearly identical between substrates with identically patterned PEG islands and micropores, further confirming disruption of cell-substrate interactions as a common mechanism behind the cellular responses on these substrates. Interestingly, cell spreading and the magnitude of migration speed was significantly greater on porous membranes compared to PEG islands with identical feature size and spacing, suggesting pore edges enhanced cellular grip. These results provide a more complete picture on how porous membranes affect cells which are grown on them in an increasing number of cellular barrier and co-culture studies.

## 1. INTRODUCTION

The interest in co-culture systems, barrier models and organ-on-a-chip studies stems from their application in drug discoveries and disease models as well as understanding of cell-cell communication at tissue interfaces.^1,2^ Due to the capabilities of porous membranes in providing biologically relevant cell-cell interactions between the co-cultured cells, these membranes have received significant attention in development of biomimetic platforms to model different tissue and organs including lung, blood brain barrier, kidney, blood vessel, gut, and liver.^3–11^ While these membranes effectively support cell-cell communication, their role in cellular behaviors including cell migration, extracellular matrix (ECM) development, and cell adhesion is not fully understood and is currently under study.

The role of porous membranes in the change of migratory behavior and ECM formation for human umbilical vein endothelial cells (HUVECs) was previously studied by our group, and it was shown that substrate disruption can lead to production of shorter fibrils and faster cell migration on porous membranes as compared to non-porous controls.^12,13^ It was concluded that the inverse relationship between fibronectin fibrillogenesis and speed of migration suggests that development of shorter fibrils on a disrupted surface can increase cell migration, or faster migration can lead to production of shorter fibrils. Although the enabling mechanism for these findings is not fully understood, it was previously suggested that the pore openings which create the surface disruption play a critical role in the regulation of these behaviors.

Still there is an unanswered question of whether surface disruption is the only contributing factor in altering migratory behavior of cells on a porous membrane, or whether pore edges should also be considered as a determinative factor for regulation of cell migration and spreading on a porous membrane. Pore edges which can provide gripping sites for cell adhesion and migration has not been previously investigated as a separate contributing factor in studies involving porous membranes. Additionally, there is a growing interest in utilizing microposts to mimic cellular microenvironment in terms of mechanical cues. These studies demonstrate how flexibility of microposts can regulate cell spreading and migratory behavior of cells through modulating substrate rigidity.^14–18^ However, similar to the role of pores in porous membranes, pillar walls can be considered as gripping points for cells, and this topological aspect of microposts can influence cell migration and spreading in addition to the substrate rigidity effect. Therefore, it is required to study cell gripping as an alternative mechanism for the aforementioned changes in cellular behavior, particularly on porous membranes which are used in an ever increasing number of scientific studies.

The aim of this study was to decouple the effect of cell gripping and surface disruption through a non-fouling micropatterned substrate. We generated a non-fouling micropattern on an SiO_2_ substrate, which resembles pores in a porous membrane in terms of shape, size and pattern of disruption, but without any pore edges. In order to create the non-fouling regions in the patterned substrate, Poly(L-lysine)-g-poly(ethylene glycol) (PLL-g-PEG) was utilized, which has been shown as a suitable candidate for generation of such non-fouling patterns in previous studies.^19–26^ The PLL backbone of this polymer allows effective adsorption onto negatively charged surfaces such as SiO_2_, while PEG branches hinder cell adhesion to the coated substrate.^19,20,24^ Stamp microcontact printing and deep UV laser ablation are commonly applied chemical patterning methods for developing such a micropatterned substrate, however as for every patterning technique they have some limitations including low reproducibility due to stamp degradation and low resolution due to the incomplete polymer ablation.^20,27,36,37,28–35^ In the current work, we combined photolithography and simple surface adsorption of the PLL-g-PEG polymer to make a reproducible pattern with fewer steps and without the aforementioned complications to provide high-resolution micropatterning. The created micropatterned substrate was used to study FN fibrillogenesis, migratory behavior, and spreading of endothelial cells, and it was demonstrated that while endothelial cells showed similar trends of changes in FN fibrilogenesis and migration speed as compared to porous substrates, different cell spreading and a lower increase in migration speed were observed on these substrates. These findings suggest that cell gripping to the pore edges should be considered as another regulatory mechanism for these behavior changes in addition to the disrupted surface.

## 2. METHODS

### 2.1 Optimization of the PLL-g-PEG coating on SiO_2_ substrate

SiO_2_ substrates were prepared using conventional microfabrication techniques similar to our previous works.12,13,38 Briefly, a 300 nm layer of SiO_2_ was prepared by plasma enhanced chemical vapor deposition (PECVD). In order to achieve non-fouling substrates, PLL(20)-g(3.5)-PEG(2) (Nanosoft Biotechnology LLC) was used, in which 20 represents a PLL backbone of 20 kDa, 3.5 is the grafting ratio and 2 corresponds to the PEG side chains of 2 kDA.^39^ To achieve high-density PLL-g-PEG grafting which effectively prevents cell adhesion, different concentrations of PLL-g-PEG in 10mM HEPES solution ranging from 0.1 to 0.9 mg/ml (0.1, 0.3, 0.5, 0.7, and 0.9 mg/ml) were placed dropwise on SiO_2_ substrate for different periods of time ranging from 30 to 115 minutes with increments of 15 minutes. After these periods of time, the solution was removed and the substrates were immediately washed before they dried. For quantification of PLL-g-PEG coating under each condition, 0.1 mg/ml fluorescent bovine serum albumin (f-BSA) was introduced to each substrate and was washed after 1 h. Prevention of BSA adsorption on SiO_2_ substrate was considered as a criterion for density of PLL-g-PEG grafting on the surface. The adsorbed f-BSA on the surface was quantified based on the florescent intensity of each surface using a Keyence BZ-X700 microscope (Keyence Corp. of America, MA, USA). Non-labeled BSA was also introduced on the SiO_2_ substrate and measured intensity was considered as the background intensity. The SiO_2_ surface without PLL-g-PEG coating was considered as the negative control and its measured intensity after introducing f-BSA was used for normalizing the measured values.

### 2.2 Development of non-fouling micropattern on SiO_2_ substrate

Photolithography steps were followed to obtain a non-fouling micropattern on the SiO_2_ layer with a similar pattern to our previous porous membranes.^12,13^ MicroPrime MP-P20 was utilized as adhesion promoter on top of the SiO_2_ layer. Microposit® S1813 positive photoresist was spin-coated at 3000 rpm for 45 s, followed by a soft bake step on a hotplate at 115 °C. GCA 6000-Series DSW 5X g-line stepper was used to expose the photoresist layer with a 150 mJ/cm2 exposure energy through a soda lime mask to obtain 3 μm pores in the photoresist with 6 μm center-to-center spacing in a hexagonal arrangement. The same method of lithography and the same mask was previously used for creating the pores in our porous membranes to have the highest consistency with our previous works.^12,13^ The exposed wafer was developed in MF CD-26 developer for 1 min, followed by washing with deionized water for 5 min. Hard bake was performed for 1 min at 150 °C. The optimum coating process which was obtained in the previous step was applied for PLL-g-PEG coating after lithography process. PLL-g-PEG solution in the optimum concentration was placed on the patterned photoresist for the optimum period of time. Then, the PLL-g-PEG solution was removed, the substrate was washed twice immediately, photoresist was removed using acetone, and the substrate was washed again with DI water. The non-fouling micropattern was visualized using f-BSA coating on patterned substrates using Keyence BZ-X700 microscope. It should be noted that the stability of PLL-g-PEG coating under acetone washing had been also evaluated by comparison between f-BSA adsorption on PLL-g-PEG coated substrates with and without acetone washing in the earlier step.

### 2.3 Atomic Force Microscope (AFM)

The PLL-g-PEG patterned substrates were examined using an MFP-3D AFM (Oxford Instruments). TR800PSA cantilevers (Oxford Instruments) with a spring constant of 0.15 N/m, were utilized to scan the surface in tapping mode.

Briefly, the sample was prepared, mounted onto a fluid cell chamber, and PBS were added on top the sample to prevent PEG brushes from collapsing onto the substrate. Surface topography scans were taken with a scan speed of 0.5Hz to minimize imaging artifacts over a fixed area. The heights of the PEG brushes were calculated from the height profile measured across the image.

### 2.4 Cell culture

Pooled human umbilical vein endothelial cells (HUVECs) were cultured in M200 with 2% Large Vessel Endothelial Supplement (LVES) and 1% penicillin and streptomycin. HUVECs and all of the cell culture reagents were purchased from Thermo Fisher unless stated otherwise. HUVECs were detached by TrypLE and seeded on samples in the density of 6×10^3^ cells/cm^2^. Cells were used between passages 3–5.

### 2.5 Cell adhesion and morphology on PLL-g-PEG coating

Preliminary experiments were required to verify that the developed coating method is effective, and PLL-g-PEG coating is efficiently non-fouling for cells. Since cells are able to attach to the uncoated areas of the patterned samples, HUVECs were instead seeded on fully PLL-g-PEG coated SiO_2_. HUVECs were also seeded on fully PLL-g-PEG coated SiO_2_ which were incubated at 37 °C for 24 h prior to cell seeding to evaluate the stability of the coating in cell culture condition. Uncoated SiO_2_ was used as the negative control.

Live/Dead viability assay was performed 24 h after cell seeding using calcein AM and ethidium homodimer-1 (EthD-1). Briefly, samples were washed with PBS and 10 μL of working solution containing 2 μM calcein AM and 4 μM EthD-1 in the cell culture medium was added to each sample. Working solution was removed after 15 min and 10 μL of cell culture medium was added to each sample, and since samples were on nontransparent Si wafer, they were inverted on the glass coverslips for fluorescence imaging. Silicone spacers were used to avoid direct contact between the samples and the glass coverslips. Images were obtained at 40X magnification with a Keyence BZ-X700 microscope through GFP and Texas RED filters for visualization of calcein AM and EthD-1, respectively.

### 2.6 Cell spreading and F-actin formation

Cells were seeded on the PEG islands and control SiO_2_ substrates for 24 h, then fixed in 3.7% Formaldehyde for 15 min, and washed with PBS. This was followed by cell permeabilization for 3 min in 0.1% Triton X-100 and washing with double-distilled water. To visualize nuclei and stress fibers, HUVECs were stained using DAPI (300 nM) and 1:400 AlexaFluor 488 conjugated phalloidin for 3 and 15 minutes, respectively. Cells were washed and 10 μL PBS was added to each sample, and the samples were flipped on a glass coverslip. Images were captured at 40X magnification through DAPI and GFP filters on a Keyence BZ-X700 microscope. Area of spreading and intensity of actin fibers were measured for each cell by ImageJ software. Intensity of actin fibers were normalized by subtracting the background intensity.

### 2.7 Fibronectin fibrillogenesis

Fibronectin fibrillogenesis was also evaluated on the PEG islands and control SiO_2_ samples. After cell culture for 24 h, cells were fixed with 3.7% formaldehyde for 15 min and washed with PBS, then blocked with 20 mg/ml BSA for 15 min and again washed with PBS. In the next step, cells were stained using 1:100 dilution of AlexaFluor 488 conjugated anti-fibronectin, Clone FN-3 for 2 h and washed with PBS three times. PBS were added to the samples and they were flipped on glass coverslips. Images were obtained at 40X magnification through a GFP filter on a Leica DMI6000 microscope (Leica Microsystems, Buffalo Grove, IL), and the lengths of fibrils were measured using a custom-written MATLAB algorithm (freely available on GitHub: https://github.com/gaborskilab). Briefly described, a disk filter with a 10-pixel radius was applied to the original image to create a background image, which was then subtracted from the original image to correct for non-uniform lighting. A Laplacian of Gaussian (LoG) filter was then applied to the background-corrected image to identify the edge of each fibronectin fibril. The half perimeter of the identified edge was reported as the fibril length.

### 2.8 Migration Assay

HUVECs were seeded on the PEG islands and control SiO_2_ substrates for 3 h and then flooded with cell culture medium containing 1:1000 dilution of SiR-DNA (Cytoskeleton Inc). An additional hour was required for cells to uptake SiR-DNA to be trackable through SiR-DNA probe using Leica DMI6000 microscope. Then, time-lapse imaging was performed every 15 minutes for 24 hours to track cell migration as described in our previous works.^13,40^ 97 frames were captured from each set stage and migration was evaluated using a custom-written MATLAB algorithm (freely available on GitHub: https://github.com/gaborskilab). Briefly, a disk filter with a 10-pixel radius was applied to the original image to create a background image, which was then subtracted from the original image to correct for non-uniform lighting. The background-corrected image was then binarized into black and white based on fluorescent intensity (any pixel that had fluorescent intensities greater than one standard deviation above the mean was considered to be part of a cell nucleus). The centroid positions of the cell nucleus from two consecutive time points defined each “step” of cell migration. The direction of each step takes on a value between 0 and 360°. The average speed of each cell was obtained by taking the ratio of the total distance traveled and the total duration of travel.

### 2.9 Statistical analysis

For statistical analysis, two-tailed student‘s t-test was used for data with Gaussian distributions and Mann–Whitney test was performed for the rest of the data. For box plots, each box includes data from 25th to 75th percentile, and the middle line shows median. The whiskers are extended to the lowest/highest values within minus/plus 1.5 times interquartile range (IQR). Sample sizes are noted in each figure legend.

## 3. RESULTS AND DISCUSSION

### 3.1 Development of a Non-Fouling Micropattern on SiO_2_ Substrates

In order to generate a non-fouling micropatterned substrate, we implemented a fabrication process utilizing PLL-g-PEG grafting on SiO_2_ substrates as shown in Figure 1. Prior to starting the patterning process, we set out to determine the highest PLL-g-PEG grafting density to achieve an effective non-fouling substrate. We varied PLL-g-PEG concentration and incubation time and determined the optimal protocol based on minimum f-BSA adsorption (Figure S1a). Very high concentrations of the polymer lowered protein repulsion. This phenomenon can be attributed to the fact that in higher concentrations, chain entanglement and lower availability of adsorption sites per polymeric chain lead to a decrease in the adsorbed segment of a polymer chain and chain loop formation instead of adopting a flat conformation on the substrate. This in turn results in easier desorption and lower polymer density on the surface.^41–44^ Since patterning of PLL-g-PEG required brief acetone washing to remove photoresist (Figure 1a), the resistance of PLL-g-PEG coating to a 3 second exposure of acetone was evaluated. The acetone exposure did not result in a reduction in the non-fouling properties of PLL-g-PEG (Figure S1b), enabling us to move forward with the patterning process.

**Figure 1.**
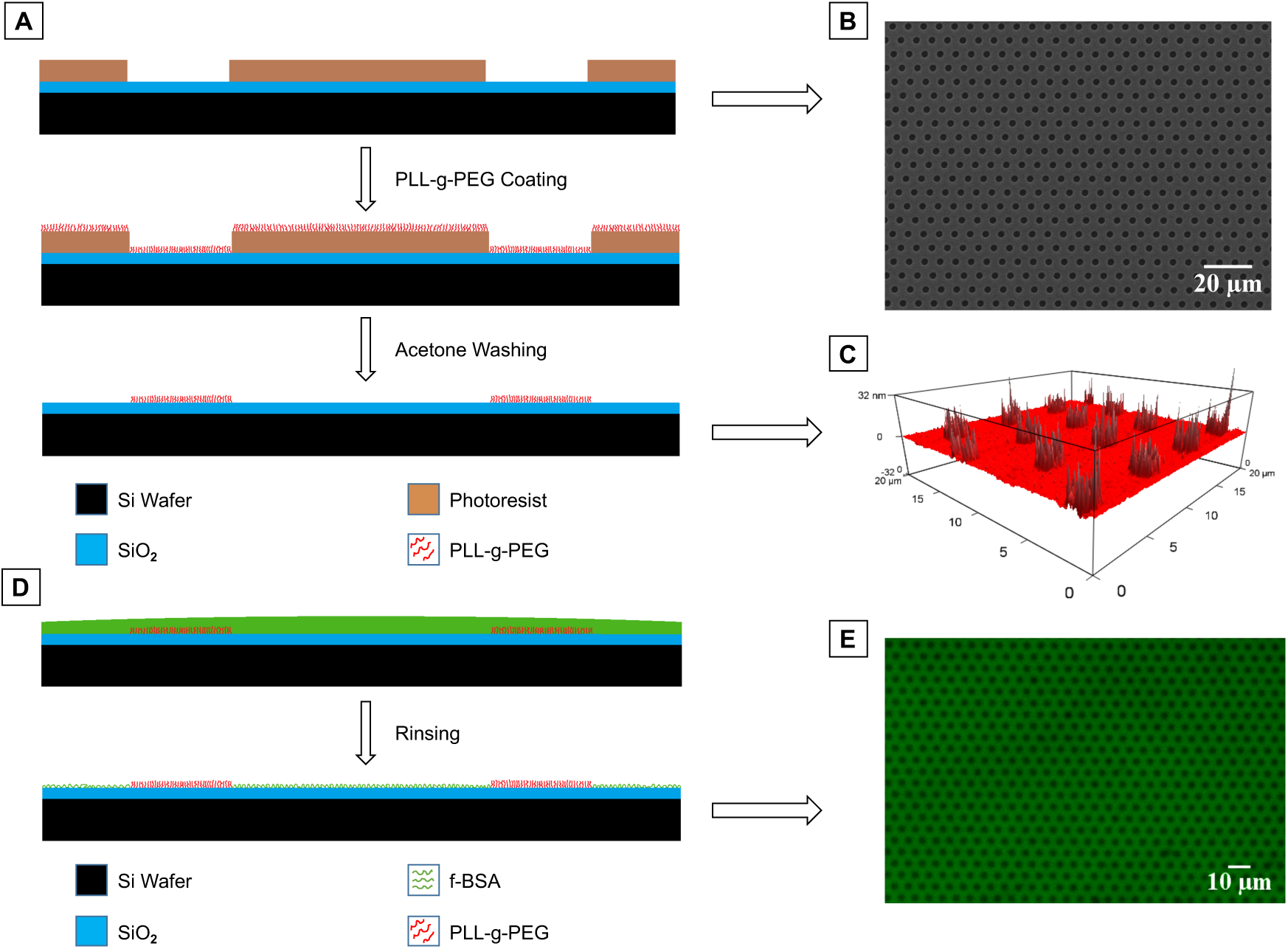
(A) Schematic illustration of preparation steps for PLL-g-PEG patterning. (B) SEM image of the surface after photolithography. (C) AFM height profile of the PLL-g-PEG-patterned substrate. (D) Process design of patterning visualization using f-BSA. (E)Visualization of the final non-fouling micropattern using f-BSA.

The developed coating protocol in the prior step was employed in combination with a photolithography process to generate the non-fouling micropattern. Two commonly used method for developing such a PLL-g-PEG pattern on our intended substrate are PDMS microcontact printing and deep UV laser ablation.^20,27,34,45^ Although microcontact printing is known to be simple and cost-effective, softness of the PDMS stamp which is necessary for suitable contact with the substrate results in vertical and lateral deformations which can affect the printed features and lead to stamp degradation after several patterning processes, particularly at our needed resolution.^30–32^ In addition, two-step ink transfer first to the stamp and then to the substrate reduces transfer efficiency and consistency of the pattern density and increases pattern defects specially when dealing with a comb co-polymer which only can be adsorbed from the backbone.^30,33^ Both mentioned complications can reduce reproducibility of the method for PLL-g-PEG patterning. On the other hand, laser ablation can provide a uniform reproducible coating on coated regions.^20,34,35^ However, determining the threshold for complete removal of the polymer without damaging the substrate can be challenging, and uncoated areas can also be covered by a low density of the patterning molecules with an incomplete ablation, which can decrease the resolution of the generated pattern.^28,29,36,37^ Therefore, we combined simple polymer surface adsorption and photolithography to make a reproducible pattern with high resolution in fewer steps. The patterning process results in a reliable production of PEG islands across the substrate. Environmental AFM measurements verified that the hydrated PLL-g-PEG molecules completely cover the patterned 3 μm regions with minimal defects (Figure 1c and S2). The limited height of the hydrated PEG molecules confirmed the process successfully resulted in a monolayer of PLL-g-PEG on the surface. The non-fouling nature of the PEG islands was visualized by incubating the substrate with f-BSA for 1 h and then washing (Figure 1d,e). The high-contrast green background with a regular pattern of black spots indicates f-BSA adsorbed to the SiO_2_ substrate everywhere except for the PEG islands. Multiple low magnification fluorescence images were taken in order to confirm faithful reproduction of the non-fouling pattern across large regions of the substrate.

### 3.2 Cell Adhesion and Morphology on Continuous PLL-g-PEG Coating

Our goal was to mimic the open pores in a membrane using patterned PEG islands. To this end, it was necessary to confirm that our method for PLL-g-PEG coating provides a non-fouling surface not just for proteins, but also cells. In order to determine if the PEG coating could prevent cell attachment, we tested uniformly coated PLL-g-PEG substrates. The substrates were prepared using the same process as for patterned islands including acetone washing, but without photoresist. Cell culture on the uniform PEG substrates verified that this coating effectively prevents cell adhesion. HUVECs were seeded on 4×4 mm samples at a density of 6000 cells/cm^2^ and imaged after 24 h. While there was a slight increase in the number of cells counted on the control SiO_2_ substrates, almost no cells were found on the PEG coated substrates (Figure 2). The very limited number of cells found on the PEG coated substrates were always rounded with limited spreading compared to cells on the uncoated substrate (Figure 2a,b). Due to concerns about long-term stability of the PLL-g-PEG coating, we pre-incubated samples for 24 h in cell culture media at 37°C prior to seeding cells.^46,47^ We found there was not a significant increase in the number of cells found on pre-incubated samples, indicating the PLL-g-PEG coating is stable for the duration of our experiments. These data confirm that the PLL-g-PEG coating reliably repels cell adhesion, mimicking the disruptive nature of pore openings.

**Figure 2.**
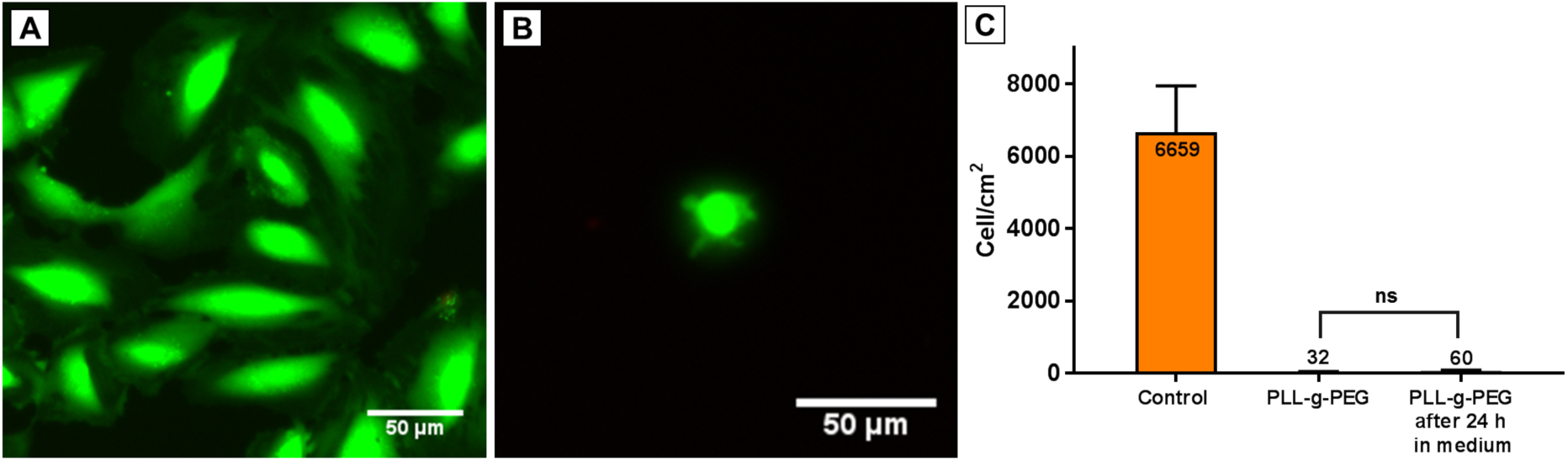
Live/Dead staining on (A) Control SiO_2_, and (B) fully PLL-g-PEG coated SiO_2_ after 24 h of cell seeding. (C) Cell density on the control SiO_2_, the fully PLL-g-PEG coated SiO_2_, and the incubated fully PLL-g-PEG coated SiO_2_ after 24 h of cell culture. (n > 7 independent substrates for each condition)

### 3.3 Cell Spreading and F-actin Quantification

Cell spreading is a result of cell-substrate interactions, and is regulated by a variety of cellular processes.^48–51^ Actin stress fibers are ubiquitous in cell adhesion and mechanotransduction, and well-organized actin fibers correspond with successful cell adhesion.^48,51,52^ We evaluated cell adhesion on the non-fouling micropatterned substrates by investigating cell spreading and stress fiber formation. HUVECs were seeded on patterned substrates, and cells were stained using DAPI and Phalloidin after 24 h to visualize cell nuclei and stress fibers, respectively. Cell spreading was measured by determining borders of each cell manually in ImageJ software. Figure 3c shows cell size distribution after 24 h of spreading on the patterned and unpatterned SiO_2_ substrates. While mean cell area on SiO2 is around 1250 μm^2^, this value was drastically lower on PEG patterned samples. This cell size reduction can originate from the frequent disruption on the SiO_2_ substrate by PLL-g-PEG islands. However, it was shown that the same pattern of disruption on SiO_2_ membranes by pores did not lead to a reduction in cell spreading.^12^ Mean fluorescent intensity of stained cytoskeleton in each cell was also measured using ImageJ to quantify stress fiber production on the patterned and unpatterned substrates. It was also found that HUVECs form fewer actin fibers on the disrupted surface of patterned (Figure 3d). The reduced cell spreading on the non-fouling micropatterns with the same surface disruption as our previous study suggests an additional factor must support cell spreading on porous membranes.

**Figure 3.**
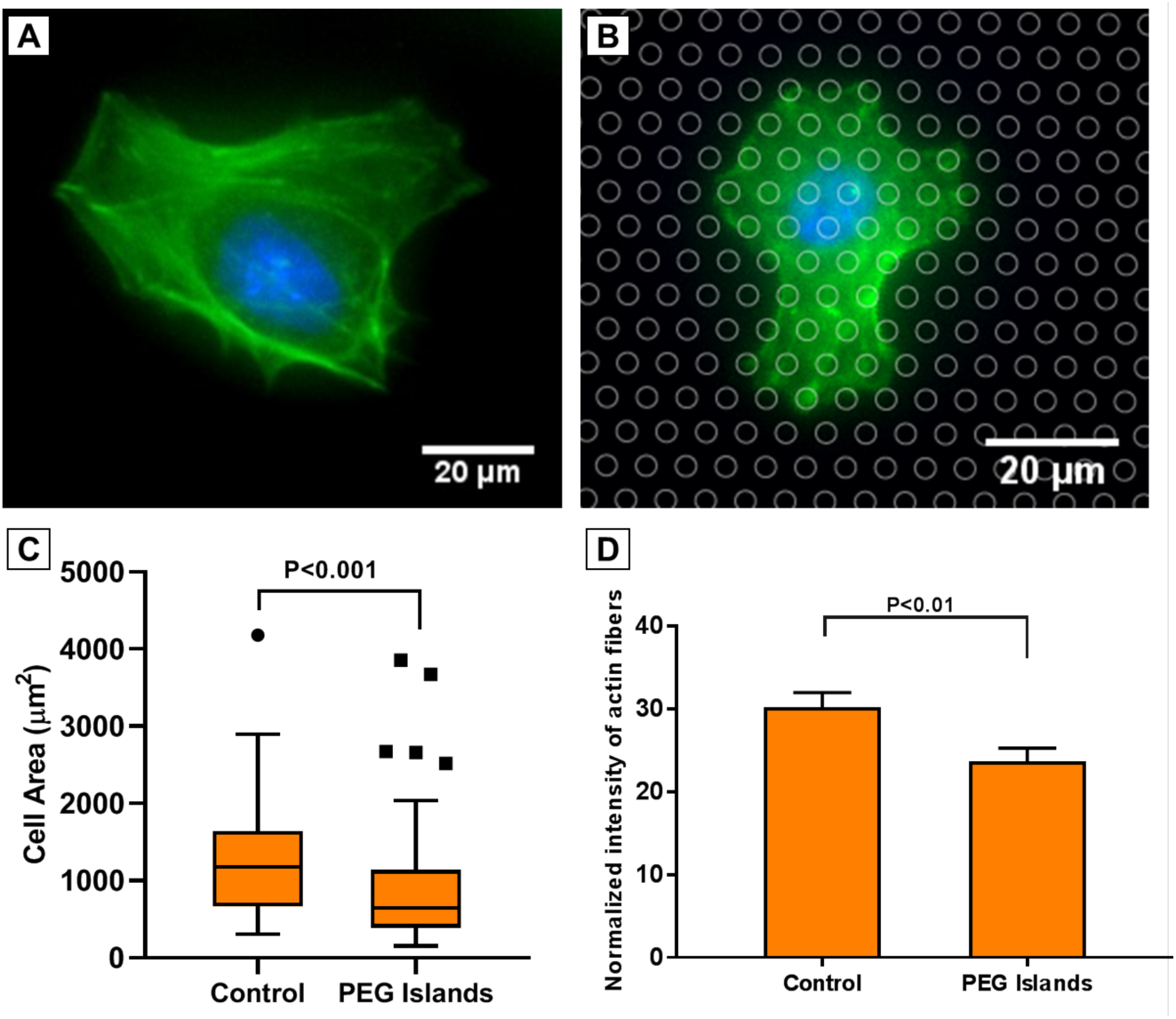
Representative images of nuclei (DAPI, blue), and F-actin (phalloidin, green) after 24h on (A) Control SiO_2_, and (B) PEG Islands. PEG islands are indicated by circles on the image B. (C, D) Cell area and Normalized intensity of F-actin, respectively. (n > 11 substrates; > 77 cells for each condition)

### 3.4 Cell Migration

To study how substrate disruption affects cell motility and directionality and whether an additional factor is at play, we compared HUVEC migration on PEG patterned and unpatterned SiO_2_ substrates. Similarly to prior studies, we investigated not only cell speed, but also the orientation or direction of cell migration between time-lapse images collected every 15 minutes over 24 h using a custom-written MATLAB algorithm (freely available on GitHub: https://github.com/gaborskilab). In these experiments, we used a far-red nuclear stain (SiR-Hoechst) that we have previously shown to have no effect on migratory behavior over long-term time-lapse imaging.^40^ As was noted earlier, cells on PEG patterned substrates spread significantly less than on control substrates. During time-lapse imaging it was also noted that some cells did not appear to fully attach and were incapable of migrating. Poorly adhered cells that did not migrate beyond a typical body width (40 μm) within 24 h were excluded from further quantitative migratory analysis. This exclusion was not necessary in our prior studies on microporous membranes and again indicated that the disruption due to PEG islands, although similar to micropore openings, resulted in some distinctly different cellular behavior.

The effect of substrate disruption on cell migration due to the PEG islands is most distinct when observing migration direction between time-lapse images. We defined each step of a cell‘s migratory trajectory based on the centroid of the nucleus from two successive images. The direction of these migration vectors indicated clear cellular preferences. We previously identified preferred sites of substrate interaction on the hexagonally micropatterened surface and labeled them as either A or B regions (Figure 4a), with three axes of symmetry. The cell steps in this study often aligned with the orientation of the A region, which closely resembles the cellular behavior on microporous substrates reported previously (Figure 4b).^13^

**Figure 4.**
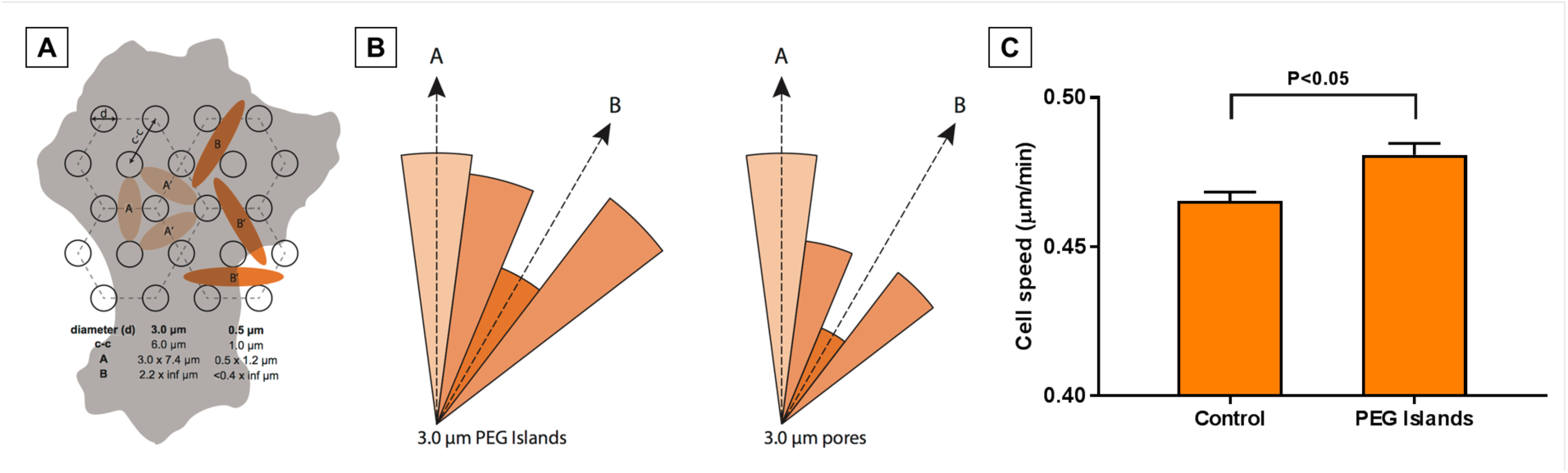
(A) Schematic illustration of the PEG island spacing and potential regions for fibronectin fibril initiations. (B) Radial histogram of the step directions for cell migration on the substrates with 3 μm PEG islands vs. 3 μm pores. (n > 400 cells; > 12000 steps) (C) Speed of migration on PEG islands vs. Control SiO_2_. (n = 4 substrates for each condition; > 1000 cells)

Using the same time-lapse images, we measured cell speed over 24 h on both PEG patterned and control substrates. Mean cell migration speed on the patterned substrates was marginally greater than that on the unpatterned substrates (Figure 4c). Cells on microporous SiO_2_ substrates also migrated faster than those on non-porous controls in our previous studies, but the differences were more striking.^13^ Taken together with the orientation data, it is clear that PEG patterning affects cell migration similarly to micropores. However, the impact of the PEG patterning on both orientation and cell speed compared to controls was less significant than that of micropores. This implies that cell-substrate disruption due to PEG islands and pore openings was only one factor affecting migratory behavior.

### 3.5 Fibronectin Fibrillogenesis

Our previous studies concluded that the discontinuity of microporous membranes disrupted cell-substrate interactions, negatively impacting FN fibrillogenesis.^12,13^ Others have described that FN fibrils preferentially form when tension is generated along a FN-substrate tether.^53^ We hypothesized that PEG islands would replicate the disruption due to the micropore openings. Here, like in our previous studies, we investigated fibronectin fibrillogenesis after 24 h on PEG patterned and unpatterned SiO_2_ substrates. Substrates were uncoated, with the only source of FN being that secreted by the cells or soluble FN in the culture media. Lengths of fibrils were measured using a custom-written MATLAB algorithm (freely available on GitHub: https://github.com/gaborskilab). Similarly to microporous substrates^12,13^, fibronectin fibril length was significantly shorter on PEG patterned SiO_2_ with a mean length of 3.2 μm compared to 5.9 μm on unpatterned SiO_2_ (Figure 5).

**Figure 5.**
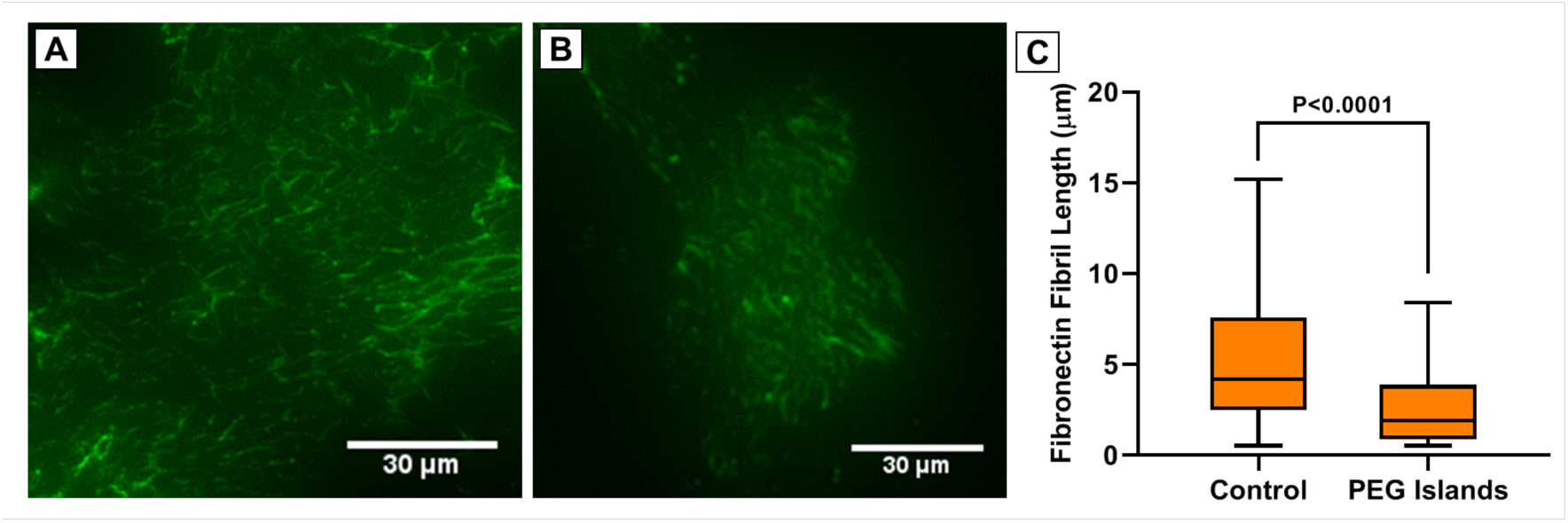
Representative images of fibronectin fibrillogenesis on (A) Control SiO_2_, and (B) PEG Islands. (C) Distribution of fibronectin fibrils lengths on Control SiO_2_, and PEG islands. (n > 5 substrates for each condition; > 3600 fibers)

### 3.6 Comparison Between PEG-Patterned and Porous SiO_2_ Substrates

Cellular behavior on our non-fouling PEG micropattern had some similarities, but several distinct differences from behavior on membranes with micropores of identical size and placement. Our intent was to recreate the cell-substrate disruptions resulting from the physical openings or void spaces of a microporous membrane. We deposited a dense and complete carpet of PLL-g-PEG within a micropattern that selectively prevented protein adsorption and therefore cell attachment within the targeted pore-like regions. The similarities in cellular behavior confirm that disruption in cell-substrate interactions is likely a major factor in the cell‘s response to a porous membrane. Some of the differences in behavior, including differences in magnitude, suggests there is another factor that contributes to cellular behavior over micropores. Due to thinness and non-fouling nature of the PEG coating (Figure 1), we expect that PLL-g-PEG does not provide any topographical cues to the cell. Micropores on the other hand, have distinct edges that the cell likely feels as it adheres and migrates over a substrate. The pore edges and walls have the same physical and chemical properties as the planar surface and may offer additional substrate interactions leading to an enhanced grip of the substrate.

We expected that discontinuity in the cell-substrate interaction through micropatterning of non-fouling PEG islands would weaken cell attachment and spreading. Indeed, we report roughly 30% reduction in HUVEC spread area over PEG islands compared to unpatterned controls (Figure 6a). Interestingly, spread area did not decrease on microporous membranes. This indicates that some aspect of the pores presents a positive factor that negates the loss in cell area due to surface discontinuity. The potential enhanced grip and additional surface area offered by the pore walls is a likely explanation of this behavior. During investigation of cell motility, speed was only marginally faster over PEG islands compared to controls, whereas microporous substrates offered a nearly 20% increase in cell speed (Figure 6b). The difference in magnitude of migration speed further points to the idea that pore edges provide enhanced cellular grip on the substrate, while the disruption may prime the cell for motility by discouraging strong and more permanent cell-substrate adhesion.^12^

**Figure 6.**
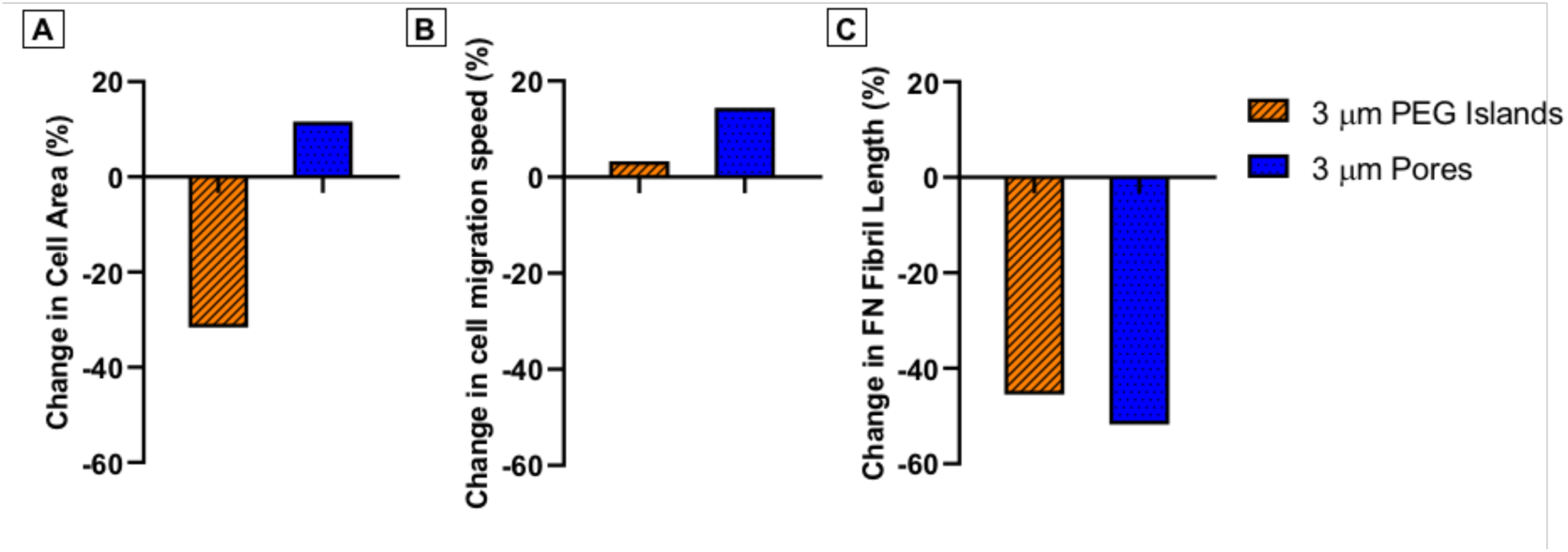
Comparative cellular behavior between 3 μm PEG islands and 3 μm pores, including (A) change in cell spread area, B) change in cell migration speed relative to unpatterned and non-porous controls, and (C) change in fibronectin fibril length.

The nearly equal reduction in FN fibril length over PEG islands and micropores indicates that the pore edges are not beneficial for fibrillogenesis. This might be surprising considering the relative increase in cell spreading and migration speed over micropores compared to PEG islands. Pore edges and walls provide increased substrate contact area, which could have offered additional FN tethering sites, and the increased spread area and cell speed suggest cells over micropores are able to generate higher traction forces and increased actin polymerization, which are positively correlated with FN fibrillogenesis.^54,55^ However, the discontinuity of a substrate due to PEG islands or micropores may dictate where fibrillogenesis occurs and dominate the response. In our previous work, a membrane with sub-micron pores had even shorter FN fibrils than the microporous membrane. The sub-micron porous membrane had less inter-pore spacing and therefore more discontinuity, but a greater number of pore edges. These data suggest that vertical pore walls may present a non-ideal placement of integrins for fibrillogenesis, but may still benefit cell spreading and migration (Figure 7).

**Figure 7.**
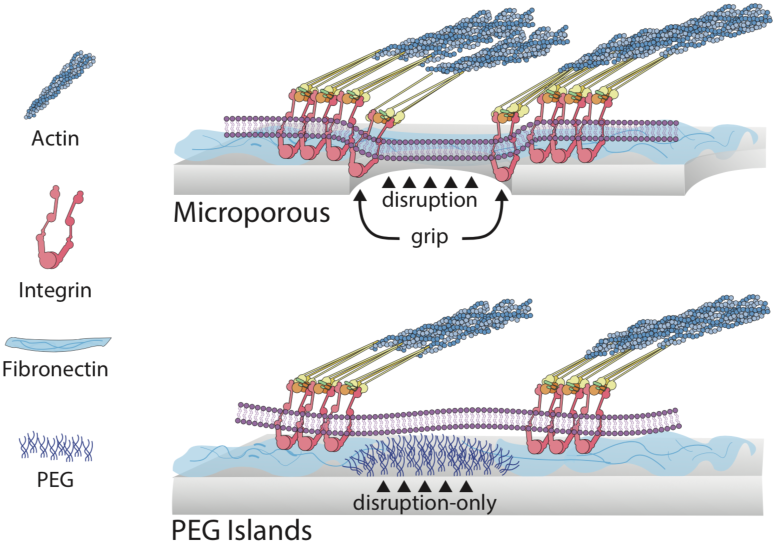
Illustration showing PEG islands disrupt cell-substrate interactions similarly to pore openings, while porous membranes also provide potential cellular grip at pore edges.

## 4. CONCLUSIONS

This study set out to determine the impact of micron-scale contact-free domains on cell-substrate interactions. Previous work showed that pore openings on cell culture membranes had significant impact on cellular behavior, including substrate adhesion, ECM production, and migration.^12,13,56^ We hypothesized that the altered behavior may be due to more than just the cell-substrate disruptions over the pore openings. Here we patterned SiO_2_ substrates with micron-scale non-fouling PEG islands that mimicked pore openings, with identical size and pattern to previously studied porous membranes. We found that while some cell-substrate interactions were disrupted similarly on patterned and porous substrates, there were also clear differences. Cell spreading and migration were enhanced on porous membranes compared to patterned PEG islands. These data suggest that membrane pores not only disrupt cell-substrate interactions, but also provide enhanced grip over non-fouling domains. This enhanced grip on the vertical pore walls, however, did not benefit FN fibrillogenesis. These results provide a more complete picture on how porous membranes affect cells which are grown on them in an increasing number of cellular barrier and co-culture studies. Future work could explore how these properties may be optimized to affect desired cellular responses to better mimic physiological conditions in these studies.

## ASSOCIATED CONTENT

Supporting Information. Optimization of PLL-g-PEG concentration and coating time for coating protocol, confirmation of PLL-g-PEG coating stability under acetone washing, Large field of view AFM 2D phase image and 3D height profile of the PLL-g-PEG patterned substrate.

## AUTHOR INFORMATION

### Author Contributions

The manuscript was written through contributions of all authors. All authors have given approval to the final version of the manuscript

### Funding Sources

Research reported in this publication was supported by NIGMS of the National Institutes of Health under award number R35GM119623 to TRG

## SUPPORTING INFORMATION

**Figure S1.**
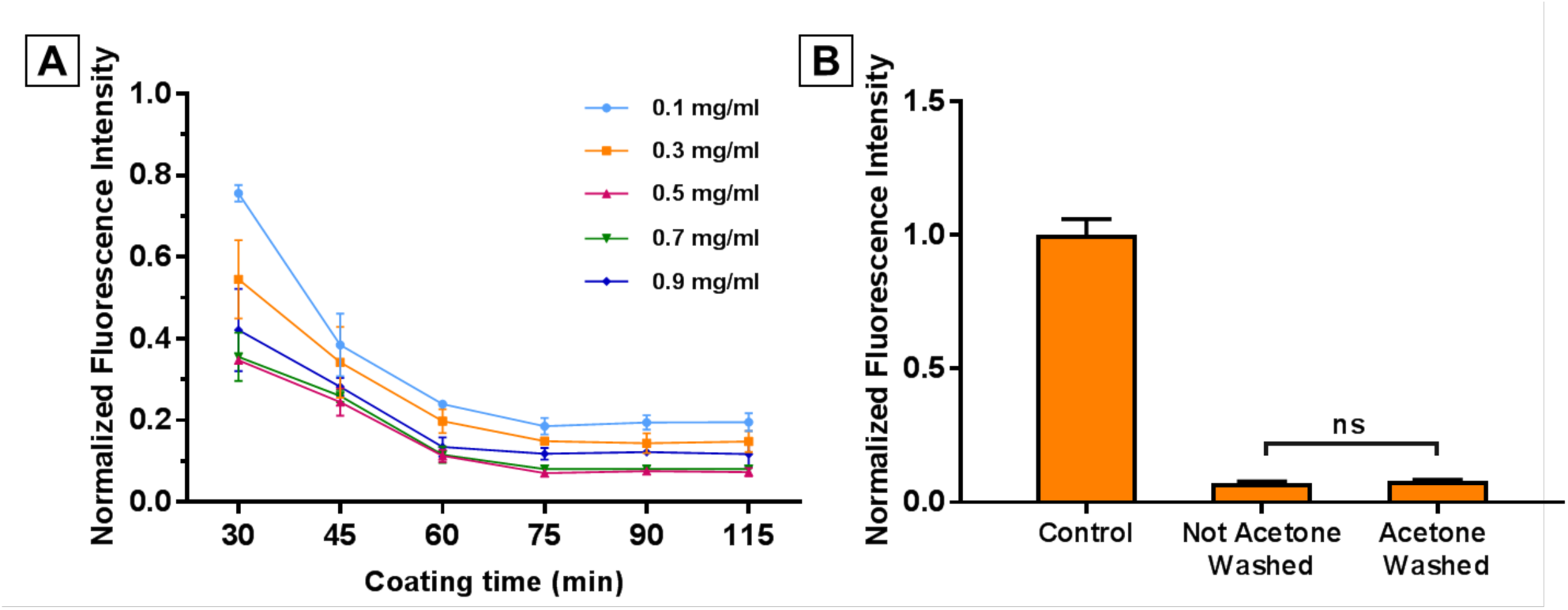
(A) Optimization of PLL-g-PEG surface functionalization protocol. Resulting adsorbed f-BSA as a function of PLL-g-PEG concentration and coating time. (B) Confirmation of PLL-g-PEG coating stability under acetone washing. For both graphs, fluorescent intensity of adsorbed f-BSA on each sample was normalized to the intensity of adsorbed f-BSA on the control untreated SiO_2_ surface. (n = 3 substrates for each condition)

**Figure S2.**
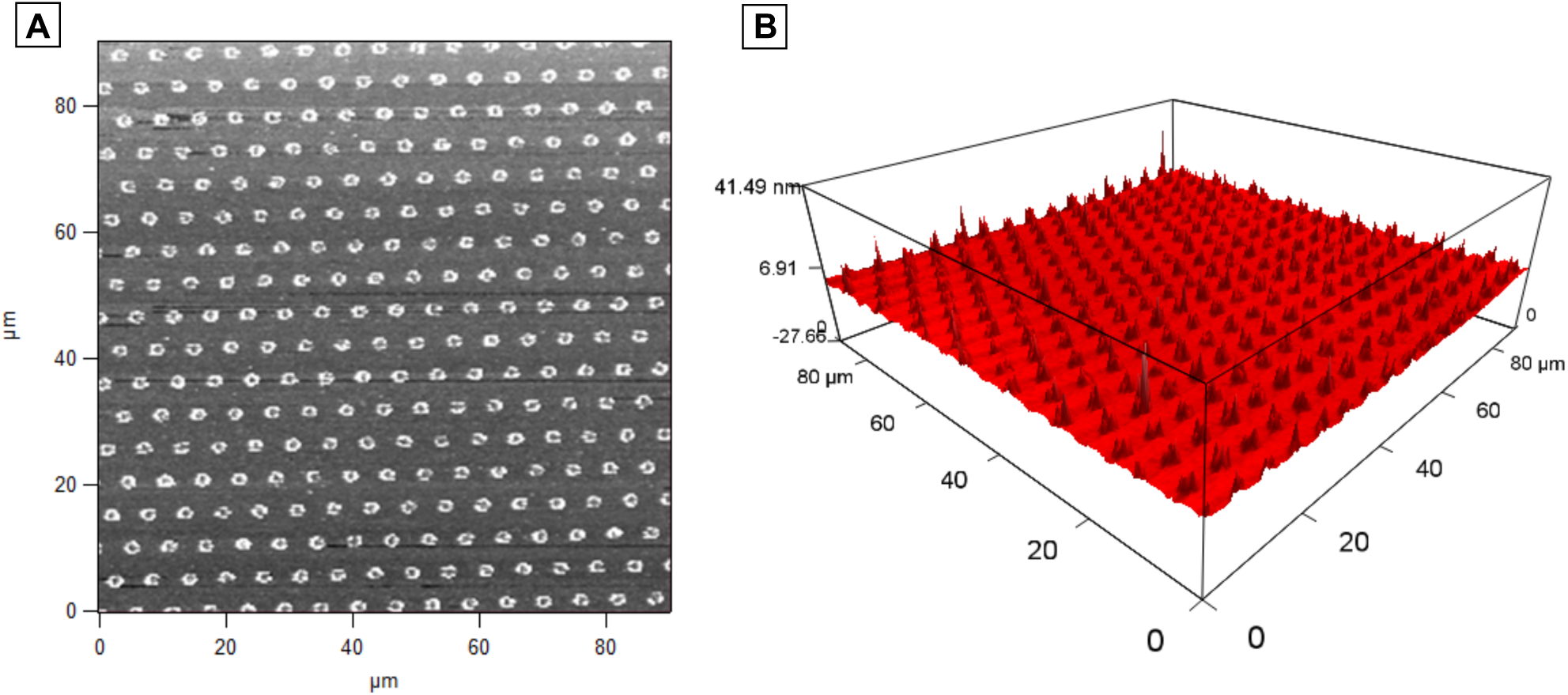
Large field of view AFM 2D phase image (A), and AFM 3D height profile (B) of the PLL-g-PEG patterned substrate under low magnification to show pattern consistency across a wide area.

## REFERENCES

(1) D. Bhatia, S. N.; Ingber, D. E. Microfluidic Organs-on-Chips. Nature Biotechnology. Nature Publishing Group August 1, 2014, pp 760–772. https://doi.org/10.1038/nbt.2989.

(2) E. Chung, H. H.; Mireles, M.; Kwarta, B. J.; Gaborski, T. R. Use of Porous Membranes in Tissue Barrier and Co-Culture Models. Lab Chip 2018, 18 (12), 1671–1689. https://doi.org/10.1039/C7LC01248A.

(3) F. Huh, D.; Matthews, B. D.; Mammoto, A.; Montoya-Zavala, M.; Hsin, H. Y.; Ingber, D. E. Reconstituting Organ-Level Lung Functions on a Chip. Science 2010, 328 (5986), 1662–1668. https://doi.org/10.1126/science.1188302.

(4) G. Booth, R.; Kim, H. Characterization of a Microfluidic in Vitro Model of the Blood-Brain Barrier (?BBB). Lab Chip 2012, 12 (10), 1784. https://doi.org/10.1039/c2lc40094d.

(5) H. Ma, S. H.; Lepak, L. A.; Hussain, R. J.; Shain, W.; Shuler, M. L. An Endothelial and Astrocyte Co-Culture Model of the Blood–Brain Barrier Utilizing an Ultra-Thin, Nanofabricated Silicon Nitride Membrane. Lab Chip 2005, 5 (1), 74–85. https://doi.org/10.1039/B405713A.

(6) I. Jang, K.-J.; Mehr, A. P.; Hamilton, G. A.; McPartlin, L. A.; Chung, S.; Suh, K.-Y.; Ingber, D. E. Human Kidney Proximal Tubule-on-a-Chip for Drug Transport and Nephrotoxicity Assessment. Integr. Biol. 2013, 5 (9), 1119–1129. https://doi.org/10.1039/c3ib40049b.

(7) J. Wang, Z.; Teoh, S. H.; Hong, M.; Luo, F.; Teo, E. Y.; Chan, J. K. Y.; Thian, E. S. Dual-Microstructured Porous, Anisotropic Film for Biomimicking of Endothelial Basement Membrane. ACS Appl. Mater. Interfaces 2015, 7 (24), 13445–13456. https://doi.org/10.1021/acsami.5b02464.

(8) K. Mazzocchi, A. R.; Man, A. J.; DesOrmeaux, J.-P. S.; Gaborski, T. R. Porous Membranes Promote Endothelial Differentiation of Adipose-Derived Stem Cells and Perivascular Interactions. Cell. Mol. Bioeng. 2014, 7 (3), 369–378. https://doi.org/10.1007/s12195-014- 0354-7.

(9) L. Pensabene, V.; Costa, L.; Terekhov, A. Y.; Gnecco, J. S.; Wikswo, J. P.; Hofmeister, W. H. Ultrathin Polymer Membranes with Patterned, Micrometric Pores for Organs-on-Chips. ACS Appl. Mater. Interfaces 2016, 8 (34), 22629–22636. https://doi.org/10.1021/acsami.6b05754.

(10) M. Kim, H. J.; Huh, D.; Hamilton, G.; Ingber, D. E. Human Gut-on-a-Chip Inhabited by Microbial Flora That Experiences Intestinal Peristalsis-like Motions and Flow. Lab Chip 2012, 12 (12), 2165. https://doi.org/10.1039/c2lc40074j.

(11) N. Granitzny, A.; Knebel, J.; Müller, M.; Braun, A.; Steinberg, P.; Dasenbrock, C.; Hansen, T. Evaluation of a Human in Vitro Hepatocyte-NPC Co-Culture Model for the Prediction of Idiosyncratic Drug-Induced Liver Injury: A Pilot Study. Toxicol. Reports 2017, 4, 89–103. https://doi.org/10.1016/J.TOXREP.2017.02.001.

(12) O. Casillo, S. M.; Peredo, A. P.; Perry, S. J.; Chung, H. H.; Gaborski, T. R. Membrane Pore Spacing Can Modulate Endothelial Cell–Substrate and Cell–Cell Interactions. ACS Biomater. Sci. Eng. 2017, 3 (3), 243–248. https://doi.org/10.1021/acsbiomaterials.7b00055.

(13) P. Chung, H. H.; Casillo, S. M.; Perry, S. J.; Gaborski, T. R. Porous Substrates Promote Endothelial Migration at the Expense of Fibronectin Fibrillogenesis. ACS Biomater. Sci. Eng. 2018, 4 (1), 222–230. https://doi.org/10.1021/acsbiomaterials.7b00792.

(14) Q. Sochol, R. D., Higa, A. T., Janairo, R. R. R., Li, S., & Lin, L. Microscale Control of Micropost Stiffness to Induce Cellular Durotaxis. In Twelfth International Conference on Miniaturized Systems for Chemistry and Life Sciences; 2008; pp 1335–1337.

(15) R. Sochol, R. D.; Higa, A. T.; Janairo, R. R. R.; Li, S.; Lin, L. Effects of Micropost Spacing and Stiffness on Cell Motility. Micro Nano Lett. 2011, 6 (5), 323. https://doi.org/10.1049/mnl.2011.0020.

(16) S. Fu, J.; Wang, Y.-K.; Yang, M. T.; Desai, R. A.; Yu, X.; Liu, Z.; Chen, C. S. Mechanical Regulation of Cell Function with Geometrically Modulated Elastomeric Substrates. Nat. Methods 2010, 7 (9), 733–736. https://doi.org/10.1038/nmeth.1487.

(17) T. Han, S. J.; Bielawski, K. S.; Ting, L. H.; Rodriguez, M. L.; Sniadecki, N. J. Decoupling Substrate Stiffness, Spread Area, and Micropost Density: A Close Spatial Relationship between Traction Forces and Focal Adhesions. Biophys. J. 2012, 103 (4), 640–648. https://doi.org/10.1016/J.BPJ.2012.07.023.

(18) U. Sochol, R. D.; Higa, A. T.; Janairo, R. R. R.; Li, S.; Lin, L. Unidirectional Mechanical Cellular Stimuli via Micropost Array Gradients. Soft Matter 2011, 7 (10), 4606. https://doi.org/10.1039/c1sm05163f.

(19) V. Falconnet, D., Pasqui, D., Park, S., Eckert, R., Schift, H., Gobrecht, J., Barbucci, J., & Textor, M. A Novel Approach to Produce Protein Nanopatterns by Combining Nanoimprint Lithography and Molecular Self-Assembly. Nano Lett. 2004, 4 (10), 1909–1914. https://doi.org/10.1021/NL0489438.

(20) W. Azioune, A.; Carpi, N.; Tseng, Q.; Théry, M.; Piel, M. Protein Micropatterns: A Direct Printing Protocol Using Deep UVs. Methods Cell Biol. 2010, 97, 133–146. https://doi.org/10.1016/S0091-679X(10)97008-8.

(21) X. Vignaud, T.; Galland, R.; Tseng, Q.; Blanchoin, L.; Colombelli, J.; Théry, M. Reprogramming Cell Shape with Laser Nano-Patterning. J. Cell Sci. 2012, 125 (Pt 9), 2134–2140. https://doi.org/10.1242/jcs.104901.

(22) Y. Rothenberg, K. E.; Neibart, S. S.; LaCroix, A. S.; Hoffman, B. D. Controlling Cell Geometry Affects the Spatial Distribution of Load Across Vinculin. Cell. Mol. Bioeng. 2015, 8 (3), 364–382. https://doi.org/10.1007/s12195-015-0404-9.

(23) Z. Liu, W.-D.; Yang, B. Patterned Surfaces for Biological Applications: A New Platform Using Two Dimensional Structures as Biomaterials. Chinese Chem. Lett. 2017, 28 (4), 675–690. https://doi.org/10.1016/J.CCLET.2016.09.004.

(24) Michel, R., Lussi, J. W., Csucs, G., Reviakine, I., Danuser, G., Ketterer, B., … & Spencer, N. D. Selective Molecular Assembly Patterning: A New Approach to Micro- and Nanochemical Patterning of Surfaces for Biological Applications. Langmuir 2002, 18 (8), 3281–3287. https://doi.org/10.1021/LA011715Y.

(25) Marie, R.; Dahlin, A. B.; Tegenfeldt, J. O.; Höök, F. Generic Surface Modification Strategy for Sensing Applications Based on Au/SiO 2 Nanostructures. Biointerphases 2007, 2 (1), 49–55. https://doi.org/10.1116/1.2717926.

(26) Lussi, J. W.; Falconnet, D.; Hubbell, J. A.; Textor, M.; Csucs, G. Pattern Stability under Cell Culture Conditions—A Comparative Study of Patterning Methods Based on PLL-g-PEG Background Passivation. Biomaterials 2006, 27 (12), 2534–2541. https://doi.org/10.1016/J.BIOMATERIALS.2005.11.027.

(27) Csucs, G.; Michel, R.; Lussi, J. W.; Textor, M.; Danuser, G. Microcontact Printing of Novel Co-Polymers in Combination with Proteins for Cell-Biological Applications. Biomaterials 2003, 24 (10), 1713–1720. https://doi.org/10.1016/S0142-9612(02)00568-9.

(28) Ihlemann, J. Patterning of Oxide Thin Films by UV-Laser Ablation. J. Optoelectron. Adv. Mater. 2005, 7 (3), 1191–1195.

(29) Musaev, O. R.; Scott, P.; Wrobel, J. M.; Wolf, J. A.; Kruger, M. B. UV Laser Ablation of Parylene Films from Gold Substrates. J. Mater. Sci. 2011, 46 (1), 183–187. https://doi.org/10.1007/s10853-010-4906-5.

(30) Falconnet, D.; Csucs, G.; Michelle Grandin, H.; Textor, M. Surface Engineering Approaches to Micropattern Surfaces for Cell-Based Assays. Biomaterials 2006, 27 (16), 3044–3063. https://doi.org/10.1016/J.BIOMATERIALS.2005.12.024.

(31) Hui, C. Y., Jagota, A., Lin, Y. Y., Kramer, E. J. Constraints on Microcontact Printing Imposed by Stamp Deformation. Langmuir 2002, 18 (4), 1394–1407. https://doi.org/10.1021/LA0113567.

(32) Huang, Y. Y., Zhou, W., Hsia, K. J., Menard, E., Park, J. U., Rogers, J. A., & Alleyne, A. G. Stamp Collapse in Soft Lithography. Langmuir 2005, 21 (17). https://doi.org/10.1021/LA0502185.

(33) Perl, A.; Reinhoudt, D. N.; Huskens, J. Microcontact Printing: Limitations and Achievements. Adv. Mater. 2009, 21 (22), 2257–2268. https://doi.org/10.1002/adma.200801864.

(34) Azioune, A.; Storch, M.; Bornens, M.; Théry, M.; Piel, M. Simple and Rapid Process for Single Cell Micro-Patterning. Lab Chip 2009, 9 (11), 1640. https://doi.org/10.1039/b821581m.

(35) Thissen, H.; Hayes, J. P.; Kingshott, P.; Johnson, G.; Harvey, E. C.; Griesser, H. J. Nanometer Thickness Laser Ablation for Spatial Control of Cell Attachment. Smart Mater. Struct. 2002, 11 (5), 792–799. https://doi.org/10.1088/0964-1726/11/5/326.

(36) Dyer, P. E. Excimer Laser Polymer Ablation: Twenty Years On. Appl. Phys. A 2003, 77 (2), 167–173. https://doi.org/10.1007/s00339-003-2137-1.

(37) Jeon, H.; Schmidt, R.; Barton, J. E.; Hwang, D. J.; Gamble, L. J.; Castner, D. G.; Grigoropoulos, C. P.; Healy, K. E. Chemical Patterning of Ultrathin Polymer Films by Direct-Write Multiphoton Lithography. J. Am. Chem. Soc. 2011, 133 (16), 6138–6141. https://doi.org/10.1021/ja200313q.

(38) Carter, R. N.; Casillo, S. M.; Mazzocchi, A. R.; DesOrmeaux, J.-P. S.; Roussie, J. A.; Gaborski, T. R. Ultrathin Transparent Membranes for Cellular Barrier and Co-Culture Models. Biofabrication 2017, 9 (1), 015019. https://doi.org/10.1088/1758-5090/aa5ba7.

(39) Müller, M.; Lee, S.; Spikes, H. A.; Spencer, N. D. The Influence of Molecular Architecture on the Macroscopic Lubrication Properties of the Brush-Like Co-Polyelectrolyte Poly(L-Lysine)-g-Poly(Ethylene Glycol) (PLL-g-PEG) Adsorbed on Oxide Surfaces. Tribol. Lett. 2003, 15 (4), 395–405. https://doi.org/10.1023/B:TRIL.0000003063.98583.bb.

(40) Chung, H. H.; Bellefeuille, S. D.; Miller, H. N.; Gaborski, T. R. Extended Live-Tracking and Quantitative Characterization of Wound Healing and Cell Migration with SiR-Hoechst. Exp. Cell Res. 2018, 373 (1–2), 198–210. https://doi.org/10.1016/J.YEXCR.2018.10.014.

(41) Lin, Y.-H. In Polymer Viscoelasticity: Basics, Molecular Theories and Experiments.; WORLD SCIENTIFIC, 2010. https://doi.org/10.1142/7786.

(42) Furusawa, K.; Yamamoto, K. Competitive Effects in Polymer Adsorption and Exchangeability of Adsorption Layer. Bull. Chem. Soc. Jpn. 1983, 56 (7), 1958–1962. https://doi.org/10.1246/bcsj.56.1958.

(43) Sperling, L. H. Polymer Surfaces and Interfaces: The Need for Uniform Terminology. ACS Div. Polym. Mater. Sci. Eng. 1995.

(44) Jiménez-Pardo, I.; van der Ven, L.; van Benthem, R.; de With, G.; Esteves, A.; Jiménez-Pardo, I.; Van der Ven, L. G. J.; Van Benthem, R. A. T. M.; De With, G.; Esteves, A. C. C. Hydrophilic Self-Replenishing Coatings with Long-Term Water Stability for Anti-Fouling Applications. Coatings 2018, 8 (5), 184. https://doi.org/10.3390/coatings8050184.

(45) Ochsner, M.; Dusseiller, M. R.; Grandin, H. M.; Luna-Morris, S.; Textor, M.; Vogel, V.; Smith, M. L. Micro-Well Arrays for 3D Shape Control and High Resolution Analysis of Single Cells. Lab Chip 2007, 7 (8), 1074. https://doi.org/10.1039/b704449f.

(46) Chen, Y.; Pidhatika, B.; von Erlach, T.; Konradi, R.; Textor, M.; Hall, H.; Lühmann, T. Comparative Assessment of the Stability of Nonfouling Poly(2-Methyl-2-Oxazoline) and Poly(Ethylene Glycol) Surface Films: An in Vitro Cell Culture Study. Biointerphases 2014, 9 (3), 031003. https://doi.org/10.1116/1.4878461.

(47) Nelson, C. M., Raghavan, S., Tan, J. L., & Chen, C. S. Degradation of Micropatterned Surfaces by Cell-Dependent and -Independent Processes†. Langmuir 2003, 19 (5), 1493–1499. https://doi.org/10.1021/LA026178B.

(48) Yeung, T.; Georges, P. C.; Flanagan, L. A.; Marg, B.; Ortiz, M.; Funaki, M.; Zahir, N.; Ming, W.; Weaver, V.; Janmey, P. A. Effects of Substrate Stiffness on Cell Morphology, Cytoskeletal Structure, and Adhesion. Cell Motil. Cytoskeleton 2005, 60 (1), 24–34. https://doi.org/10.1002/cm.20041.

(49) Discher, D. E.; Janmey, P.; Wang, Y.-L. Tissue Cells Feel and Respond to the Stiffness of Their Substrate. Science 2005, 310 (5751), 1139–1143. https://doi.org/10.1126/science.1116995.

(60) Pelham, R. J.; Wang, Y. l. Cell Locomotion and Focal Adhesions Are Regulated by Substrate Flexibility. Proc. Natl. Acad. Sci. U. S. A. 1997, 94 (25), 13661–13665. https://doi.org/10.1073/PNAS.94.25.13661.

(61) Cavalcanti-Adam, E. A.; Micoulet, A.; Blümmel, J.; Auernheimer, J.; Kessler, H.; Spatz, J. P. Lateral Spacing of Integrin Ligands Influences Cell Spreading and Focal Adhesion Assembly. Eur. J. Cell Biol. 2006, 85 (3–4), 219–224. https://doi.org/10.1016/J.EJCB.2005.09.011.

(62) Tojkander, S.; Gateva, G.; Lappalainen, P. Actin Stress Fibers--Assembly, Dynamics and Biological Roles. J. Cell Sci. 2012, 125 (Pt 8), 1855–1864. https://doi.org/10.1242/jcs.098087.

(63) Singh, P.; Carraher, C.; Schwarzbauer, J. E. Assembly of Fibronectin Extracellular Matrix. Annu. Rev. Cell Dev. Biol. 2010, 26, 397–419. https://doi.org/10.1146/annurev-cellbio-100109-104020.

(64) Lemmon, C. A.; Chen, C. S.; Romer, L. H. Cell Traction Forces Direct Fibronectin Matrix Assembly. Biophys. J. 2009, 96 (2), 729–738. https://doi.org/10.1016/J.BPJ.2008.10.009.

(65) Eisenberg, J. L.; Safi, A.; Wei, X.; Espinosa, H. D.; Budinger, G. S.; Takawira, D.; Hopkinson, S. B.; Jones, J. C. Substrate Stiffness Regulates Extracellular Matrix Deposition by Alveolar Epithelial Cells. Res. Rep. Biol. 2011, 2011 (2), 1–12. https://doi.org/10.2147/RRB.S13178.

(66) Salminen, A. T.; Zhang, J.; Madejski, G. R.; Khire, T. S.; Waugh, R. E.; McGrath, J. L.; Gaborski, T. R. Ultrathin Dual-Scale Nano- and Microporous Membranes for Vascular Transmigration Models. Small 2019, 15 (6), 1804111. https://doi.org/10.1002/smll.201804111.

